# Variation in moment-to-moment brain state engagement changes across development and contributes to individual differences in executive function

**DOI:** 10.1101/2024.09.06.611627

**Authors:** Jean Ye, Link Tejavibulya, Wei Dai, Lora M. Cope, Jillian E. Hardee, Mary M. Heitzeg, Sarah Lichenstein, Sarah W. Yip, Tobias Banaschewski, Gareth J. Baker, Arun L.W. Bokde, Rüdiger Brühl, Sylvane Desrivières, Herta Flor, Penny Gowland, Antoine Grigis, Andreas Heinz, Jean-Luc Martinot, Marie-Laure Paillère Martinot, Eric Artiges, Frauke Nees, Dimitri Papadopoulos Orfanos, Luise Poustka, Sarah Hohmann, Nathalie Holz, Christian Baeuchl, Michael N. Smolka, Nilakshi Vaidya, Henrik Walter, Robert Whelan, Gunter Schumann, Hugh Garavan, Bader Chaarani, Dylan G. Gee, Arielle Baskin-Sommers, BJ Casey, IMAGEN consortium, Dustin Scheinost

**Author notes:** Correspondence: Jean Ye, 300 Cedar St., New Haven, CT.

## Abstract

Neural variability, or variation in brain signals, facilitates dynamic brain responses to ongoing demands. This flexibility is important during development from childhood to young adulthood, a period characterized by rapid changes in experience. However, little is known about how variability in the engagement of recurring brain states changes during development. Such investigations would require the continuous assessment of multiple brain states concurrently. Here, we leverage a new computational framework to study state engagement variability (SEV) during development. A consistent pattern of SEV changing with age was identified across cross-sectional and longitudinal datasets (N>3000). SEV developmental trajectories stabilize around mid-adolescence, with timing varying by sex and brain state. SEV successfully predicts executive function (EF) in youths from an independent dataset. Worse EF is further linked to alterations in SEV development. These converging findings suggest SEV changes over development, allowing individuals to flexibly recruit various brain states to meet evolving needs.

## Introduction

Neural variability describes variation in brain signals (1). Using a range of neuroimaging modalities, studies have characterized neural variability in several ways. These include variability in dynamic functional connectivity and trial-to-trial brain BOLD signals (2,3). Once considered a source of measurement noise, neural variability is now appreciated for its ability to support executive functions (EF;2). Specifically, increased neural variability allows the exploration of different brain network combinations before adopting the most optimal one for the task at hand (2). Elevated neural variability has been consistently linked to better performance in EF tasks (1,4–6).

Given the significant neurodevelopment and EF improvement during adolescence (7–10), the role of neural variability is particularly crucial to consider during this critical life period. Perseverative behavioral patterns observed in younger children give way to the more flexible responses seen in older individuals across adolescent development (11,12). Adolescence is additionally characterized by rapid changes in experiences and increased independence (13). These changes can translate into a higher demand for flexibility to meet evolving needs. Development in neural variability may help meet these unique challenges. Indeed, previous literature has reported brain signal variability increasing with age during development (4,14). Age-related increases in functional connectivity pattern variability have been observed in developmental populations (6,15,16). However, much remains unknown about variability regarding brain states (i.e., activation or connectivity patterns recurring across time and task conditions), an increasingly popular concept in neuroimaging research (17), in the developmental population.

Recent work has examined variability in brain states by studying how adolescents transitioned from one brain state to another (5). However, it is unclear how variability in the continuous, moment-to-moment engagement of any one given brain state changes with age and supports behavioral maturation. Examining each brain state individually is important, as the developmental trajectory and behavioral relevance of their engagement variability may vary. Answering these questions would require the continuous tracking of multiple brain states simultaneously. As many brain dynamic methods tend to characterize each time unit with only the dominant brain state, they do not permit temporal overlap in brain state engagement. Thus, a moment-to-moment quantification of engagement for each brain state is unavailable, making it challenging to assess its continuous engagement. As a result, the developmental trajectory and behavioral relevance of brain state engagement variability (SEV) remain unknown.

We investigated these questions by capitalizing on a new multivariate computational framework (18) that circumvents earlier limitations. Utilizing non-negative least squares regression, this approach not only provides state engagement information at the resolution of individual time points, but also allows temporal overlap in brain state recruitment. We leveraged this framework to estimate SEV in resting-state, naturalistic, and task-based functional magnetic resonance imaging (fMRI) from two cross-sectional and two longitudinal datasets. We began by investigating whether age-related changes in SEV appear and generalize across paradigms. Sex differences in neurodevelopment exist (19). Using self-reported sex information, we explored sex-by-age interactions in SEV. Next, we examined the relationship between SEV and EF in youth. To this end, predictive models of individual differences in EF based on SEV were created using machine learning. Mediation models further tested the hypothesis that SEV would change with age to support EF. Lastly, we studied whether larger alterations (i.e., the squared difference between chronological and brain age) from typical SEV development were associated with differences in EF.

## Results

We first investigated SEV in two publicly available cross-sectional datasets, the Philadelphia Neurodevelopmental Cohort (PNC; 20) and the Healthy Brain Network (HBN; 21). Resting-state fMRI data from both datasets (PNC: N=1208, 658 females; HBN: N=1275, 491 females) were analyzed. We additionally included HBN’s naturalistic fMRI data (N=1313, 505 females), which were collected while participants watched a clip from the movie *Despicable Me*.

As both datasets were cross-sectional, we further validated our findings in two longitudinal datasets, the IMAGEN study (22,23) and the Michigan Longitudinal Study (MLS; 24,25). Task-based fMRI data collected during a cognitive control paradigm were analyzed (IMAGEN: Stop Signal; MLS: Go/NoGo). For the IMAGEN cohort, longitudinal fMRI data were collected from 530 participants (319 females) at 14, 19, and 22 years old. Two different MLS cohorts were used: one including 103 participants (40 females) aged 7.6 to 21.7 years with 479 total visits (MLS cohort 1), and the other comprising 150 participants (56 females) between 16.1 to 28.5 years of age with 639 total visits (MLS cohort 2).

To assess SEV, we first implemented our established pipeline to identify four canonical brain states using nonlinear manifold learning and task-based fMRI data from the Human Connectome Project (HCP) dataset (26,27). We used fMRI data from six tasks (motor, working memory, social, emotional, relational, and gambling). As this dataset included tasks ranging from motor to cognitive and affective paradigms, it has the potential to reveal brain states underlying a diverse set of cognitive processes. Additionally, identifying brain states in another dataset avoids circular analysis and overfitting. Based on their associated task conditions and demands, we labeled the identified brain states as fixation, high-cognition, low-cognition, and cue/transition (**Supplementary Material**). The fixation state mostly contained time points from the fixation condition. The high-cognition state included time points from complex cognitive paradigms such as working memory, emotion, relational, gambling and social. The low-cognition state included a large number of time points from the motor task. Finally, the cue/transition state consisted mostly of time points from the cue condition. The activation of canonical functional brain networks also followed what cognitive processes were associated with each state. For instance, the high-cognition state showed frontoparietal network activation and default mode network deactivation (**Supplementary Material**).

The continuous engagement of these four brain states was evaluated in the PNC, HBN, IMAGEN, and MLS cohorts (**Figure 1;** for details, see 18). SEV was operationalized as the standard deviation of moment-to-moment engagement across time.

**Figure 1.**
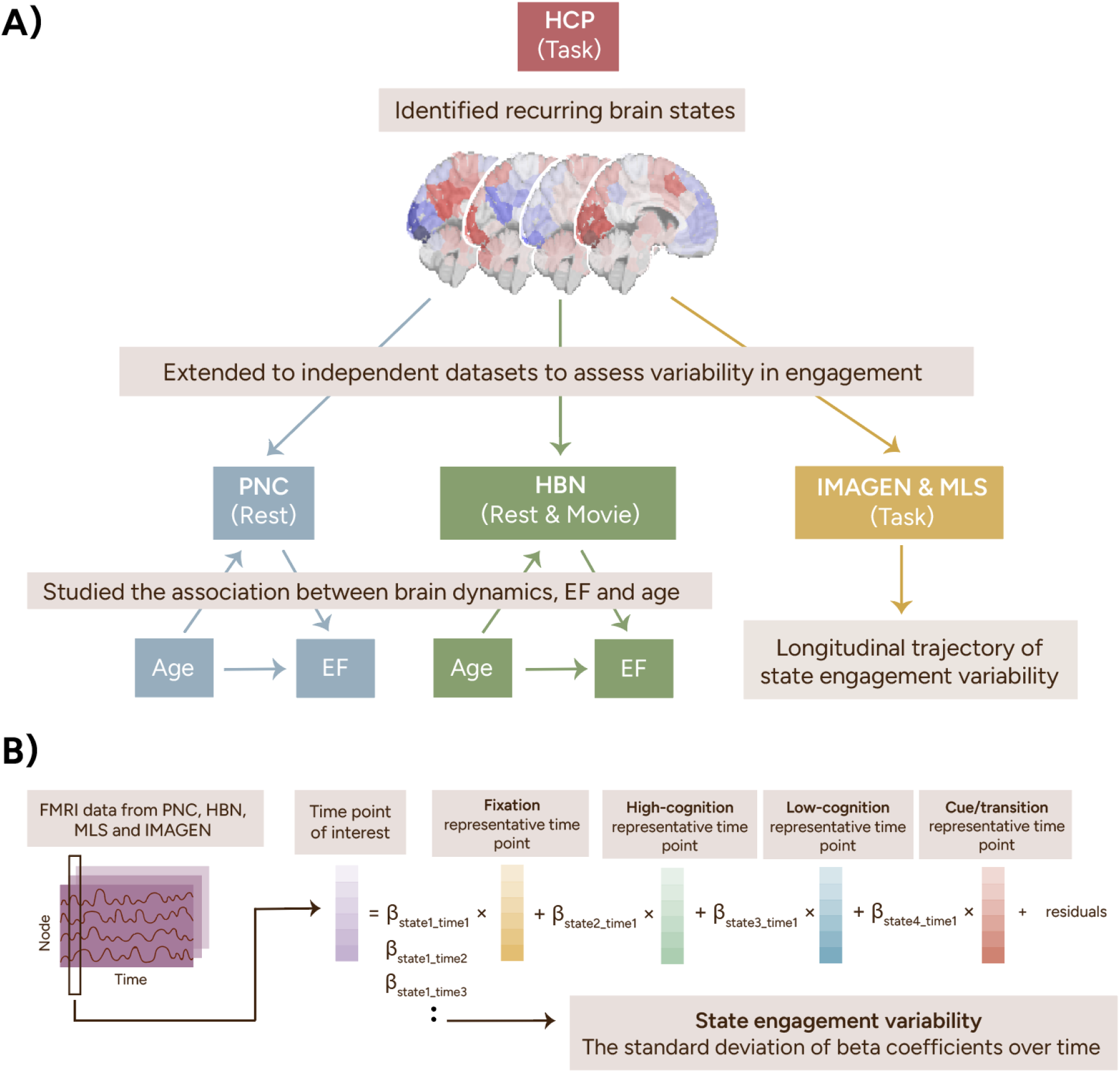
Study pipeline. We identified four recurring brain states using task-based fMRI data from the HCP. These four brain states were then extended to the PNC, HBN, IMAGEN, and MLS datasets using a previously established pipeline shown in **B**. To summarize, the representative time points from all four states were regressed from each time point from the PNC, HBN, IMAGEN, and MLS datasets using non-negative least squares regression. This step returned a beta coefficient for each state, indicating its engagement at that time point. State engagement variability was computed as each participant’s standard deviation of beta coefficients across time. HCP, Human Connectome Project; PNC, Philadelphia Neurodevelopmental Cohort; HBN, Healthy Brain Network; MLS, Michigan Longitudinal Study.

### SEV increased with age across fMRI paradigms

In both PNC and HBN, we observed a positive association between age and SEV during resting-state and movie-watching (MANOVA; PNC rest: F(4,1201)=20.793, p<0.001; HBN rest: F(4,1268)=36.482, p<0.001; HBN movie: F(4,1305)=55.959, p<0.001). Post-hoc correlations highlighted that this positive association can be found across states and cohorts (**Figure 2**). Nevertheless, these effect sizes demonstrated variations. Stronger effects were observed in HBN compared to PNC and in movie compared to rest. The high-cognition and fixation states showed the greatest effects out of all brain states examined.

**Figure 2.**
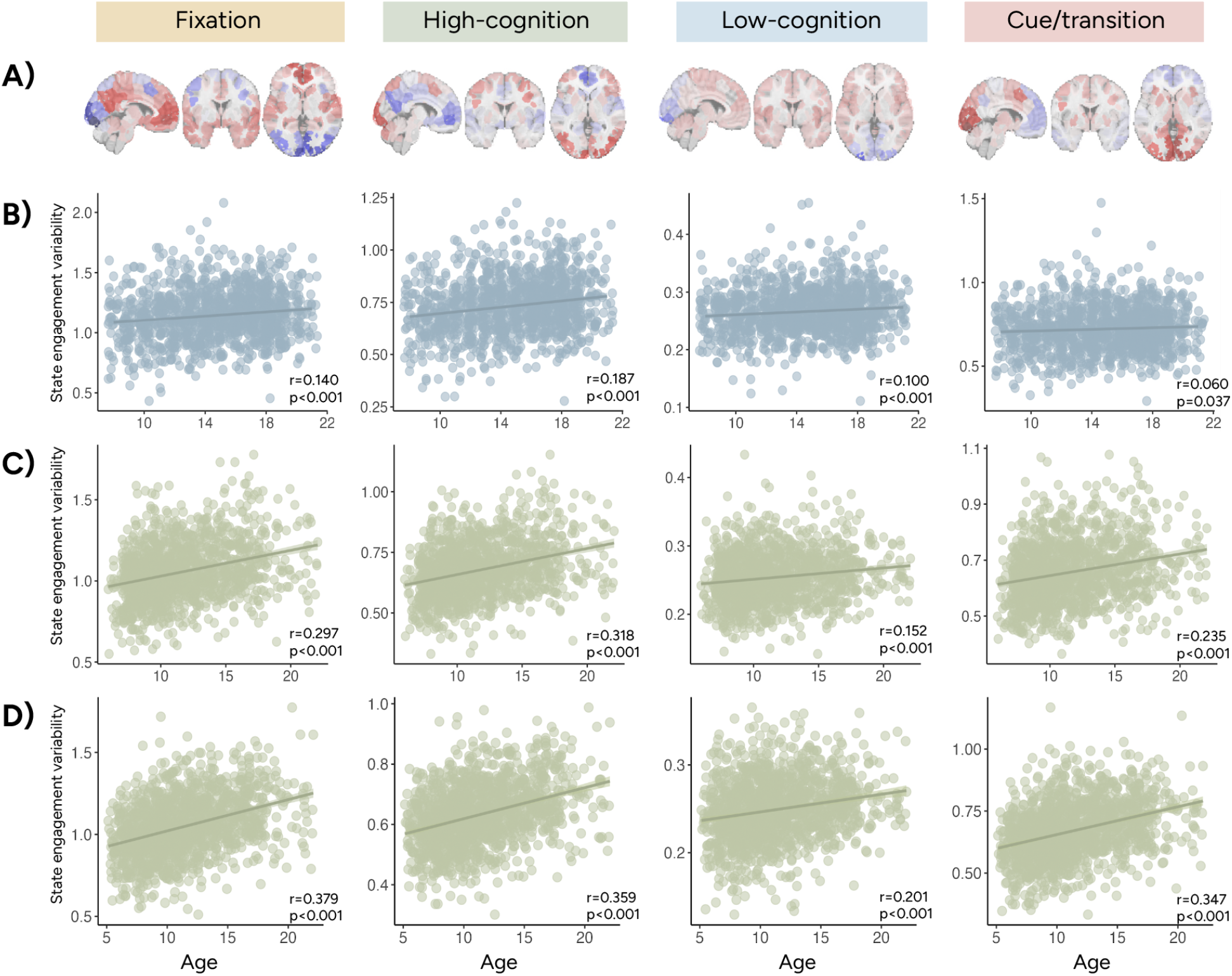
State engagement variability increased with age in PNC and HBN. **A** shows the activation patterns associated with the fixation, high-cognition, low-cognition, and cue/transition brain states. Across all four brain states, we observed a positive association between age and state engagement variability in PNC rest **(B)**, HBN rest **(C)**, and HBN movie **(D)**. Results from Pearson correlation between state engagement variability and age are also reported here.

### Longitudinal changes in SEV

As cross-sectional data might not be sensitive to subtle developmental changes, we investigated longitudinal changes in SEV by applying linear mixed-effects (LME) models to longitudinal fMRI data. Data were collected at ages 14, 19 and 22 in the IMAGEN dataset. Fixation and high-cognition SEV was greater at ages 19 and 22 compared to age 14 (**Figure 3; Supplementary Table 3**). Interestingly, no significant age difference was found for low-cognition or cue/transition SEV (**Figure 3; Supplementary Table 3**). SEV additionally did not differ between age 19 and 22 across all four states (**Supplementary Table 3**). These results extended our previous cross-sectional results, suggesting that SEV does not increase with age indefinitely, but rather stabilizes around mid-adolescence. Importantly, the timing of this asymptote may be brain state dependent.

**Figure 3.**
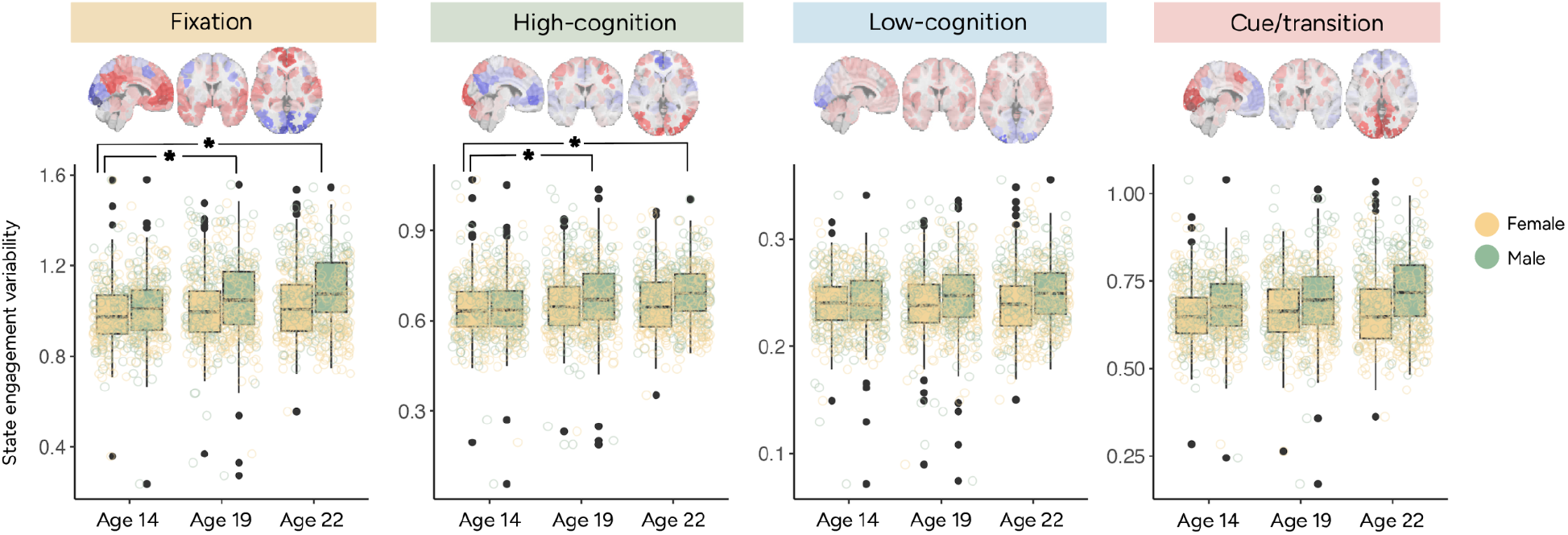
State engagement variability in the IMAGEN dataset plotted by visit and sex. Data were collected at ages 14, 19 and 22. Ages 19 and 22 showed greater fixation and high-cognition state engagement variability than those aged 14. However, no significant age difference was observed for low-cognition and cue/transition state engagement variability. Ages 19 and 22 did not differ significantly in state engagement variability for any brain states examined.

This possibility was further explored in MLS, which included multiple visits from some participants and allowed for the investigation of smoother trajectories. In the younger sample (MLS cohort 1; ages 7.6-21.7 years), SEV increased with age across all four states (fixation: beta=0.028, t-value(470.8)=5.07, p<0.001; high-cognition: beta=1.916e-02, t-value(470.7)=5.321, p<0.001; low-cognition: beta=1.972e-03, t-value(460.3)=2.002, p=0.046; cue/transition: beta=0.012, t-value(468.9)=3.670, p<0.001; **Supplementary Table 4** for other covariates; **Supplementary Figure 1**). However, consistent with our prior observations in IMAGEN, SEV did not change significantly with age in the older sample (MLS cohort 2; 16.1-28.5 years; LME; fixation: beta=-0.001, t-value(584.6)=-0.246, p=0.806; high-cognition (beta=0.0006, t-value(580.3)=0.191, p=0.849; low-cognition: beta=-0.0007, t-value(590.4)=-0.771, p=0.441; cue/transition: beta: −0.002, t-value(587.793)=-0.514, p=0.607; **Supplementary Table 5** for other covariates). To visualize this developmental trajectory, we combined both MLS cohorts. These trajectories (**Figure 4**) suggest that SEV follows a nonlinear trajectory. Observations across four independent datasets aligned to suggest a general pattern of SEV first increasing with age before stabilizing around mid-adolescence. However, specific timings of stabilization may vary by brain state.

**Figure 4.**
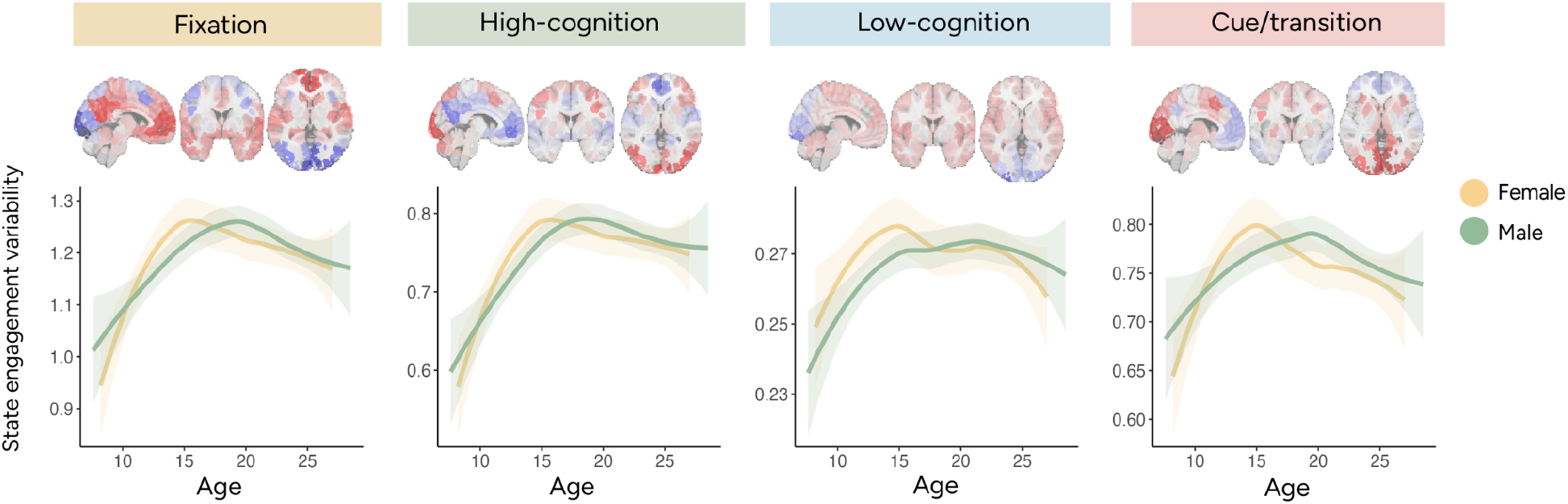
State engagement variability trajectories plotted by sex using data from the MLS datasets. The smoothed curve was generated using the Locally Weighted Least Square Regression technique in R’s ggplot2 package. Activation patterns associated with each brain state are again shown above the trajectories. Across all four brain states, state engagement variability increased with age before reaching its peak in adolescence. Plotting trajectories by group additionally indicated that female participants appeared to reach a stabilization point before male participants.

### Timing of SEV stabilization varied by self-reported sex

As previous work has noted sex differences in brain development, we further probed sex effects on SEV using self-reported sex information. A significant age-by-sex interaction was found in PNC (MANOVA; F(4,1201)=3.838, p=0.004; **Supplementary Table 6**) but not in HBN (MANOVA; rest: F(4,1268)=0.54, p=0.706; movie: F(4,1305)=0.394, p=0.813; **Supplementary Table 6**). However, age differences between the two datasets likely contributed to this discrepancy (two-sample t-tests: p<0.001; PNC age: 14.688±3.321; HBN rest age: 11.693±3.39; HBN movie: 11.361±3.606). Compared to HBN, the PNC cohort included more participants in late adolescence. The age-by-sex interaction in PNC likely reflects potential group differences occurring later in development, perhaps when individuals reach the stabilization point previously noted in our longitudinal results (**Figure 4**).

In support of this hypothesis, we observed a significant interaction between sex and later visit on SEV across all four states in IMAGEN (LME; interaction between sex and visit at 22 years old; fixation: beta=5.390e-02, t(1056)=3.429, p<0.001; high-cognition: beta=3.741e-02, t(1056)=3.609, p<0.001; low-cognition: beta=1.190e-02, t(1056)=3.928, p<0.001; cue/transition: beta=3.425e-02, t(1056)=3.496, p<0.001; see **Supplementary Table 7** for interaction between sex and visit at 19 years old). These results provide further evidence of potential sex differences occurring in late adolescence.

To further probe whether there were sex differences in the timing of when SEV peaked, we performed post-hoc analysis in the MLS dataset, which included dense sampling across various ages. The peak age was first identified for each sex (implemented using segmented.lme in R; **Table 1**), and the difference between the two sexes was calculated. Next, the peak age difference was recalculated 500 times after shuffling self-reported sex across participants. We subsequently determined the p-value by counting the number of permutation tests yielding a larger difference than the observed value.

**Table 1.** Age at transition point in state engagement variability trajectory by sex (MLS)

|  | Fixation | High-cognition | Low-cognition | Cue/transition |
| --- | --- | --- | --- | --- |
| Male | 17.5764 | 15.8559 | 13.6781 | 17.4534 |
| Female | 16.0619 | 15.8555 | 13.6784 | 16.4343 |

Permutation testing revealed that female participants arrived at the peak age earlier for the fixation brain state (p<0.001) but not for the other brain states (high-cognition: p=0.82; low-cognition: p>0.999; cue/transition: p=0.154). It is important to note here that the two groups showed more overlap in variations around the peak age for the high-cognition, low-cognition, and cue/transition brain states (**Figure 4**), potentially contributing to the lack of sex differences observed.

### SEV predicts EF in development

Our findings revealed a remarkably consistent pattern of SEV changing with age. As development is characterized by dramatic changes in experiences, one follow-up question is whether variations in SEV have any behavioral implications. We investigated the association between SEV and EF in the two cross-sectional datasets. EF was evaluated using the NIH Toolbox and the Penn Computerized Neurocognitive Battery (CNB) in HBN and PNC, respectively. A summary EF score was extracted using Principal Component Analysis (PCA) on scores from the different EF tasks (see **Methods; Supplementary Figure 2**). In the PNC dataset, the 1st EF component accounted for 48.166% of the variance. The 1st EF component accounted for 65.379% and 68.205% of the variance in the HBN rest and movie cohorts, respectively.

First, we examined whether SEV can predict EF in previously unseen individuals from an independent dataset. A linear model was trained to predict EF using SEV in either dataset before being applied to the other dataset. To keep the fMRI paradigm consistent in both the training and testing datasets, we analyzed the PNC and HBN rest cohorts. Predicted EF showed a significant positive correlation with observed EF (Pearson correlation; model tested in PNC: r=0.160, p<0.001; model tested in HBN: r=0.202, p<0.001). We successfully predicted EF with relatively few features included in the models (see model parameters in **Supplementary Table 8**), suggesting that SEV closely supports EF performance. Importantly, these consistent results were obtained from two unharmonized datasets evaluating EF utilizing different tools.

### SEV mediates the association between age and EF

As EF performance tends to improve with age in development, we next tested whether SEV mediated the relationship between age and EF (28). For this analysis, an overall SEV measure was extracted by performing PCA on the four SEV measures. The 1st SEV PCA component accounted for 92.786%, 91.657%, and 94.212% of the variance for PNC rest, HBN rest, and HBN movie, respectively. Pearson correlations were performed between variables before they were entered into a mediation model. In line with previous literature (10,29), EF scores increased with age (PNC: r=0.498, p<0.001; HBN rest: r=0.631, p<0.001; HBN movie: r=0.685, p<0.001). Overall SEV positively correlated with EF when controlling for age (PNC: r=0.089, p=0.002; HBN rest: r=0.071, p=0.015; HBN movie: r=0.12, p<0.001). Mediation models revealed that overall SEV partially mediated the relationship between EF and age across cohorts (PNC: beta=0.01, p-value=0.001; HBN rest: beta=0.13, p-value=0.018; HBN movie: beta=0.29, p-value=0.0003; direct effect reported in **Figure 5**).

**Figure 5.**
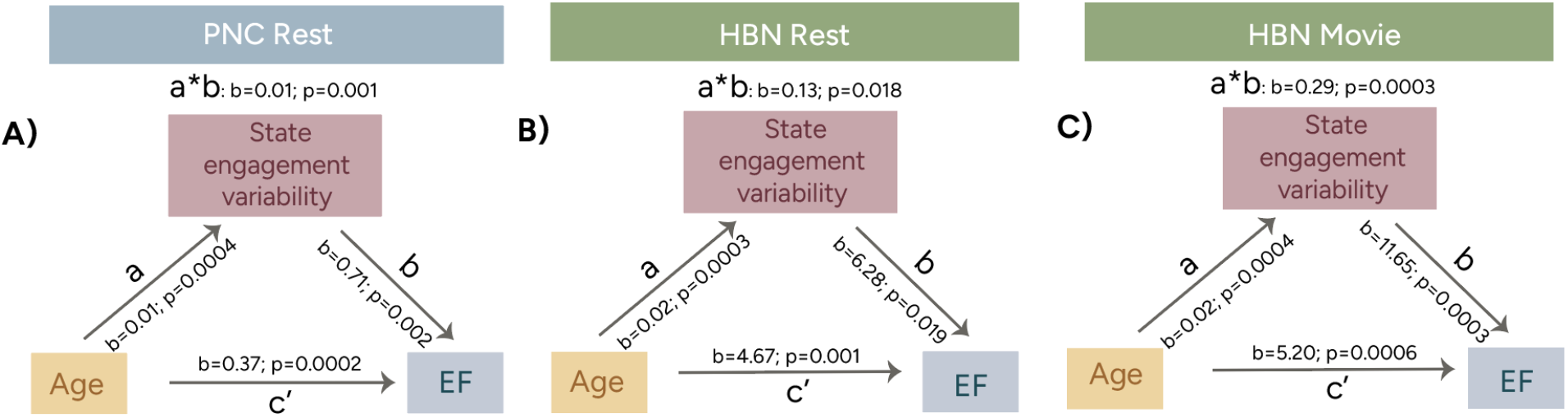
Overall state engagement variability partially mediated the relationship between age and EF in the PNC rest **A)**, HBN rest **B)** and HBN movie **C)** cohorts. a: age’s effect on state engagement variability; b: state engagement’s effect on EF; c’: the direct effect; a*b: the indirect effect. Mediation analysis was performed using the Mediation Toolbox (27).

**Figure 6.**
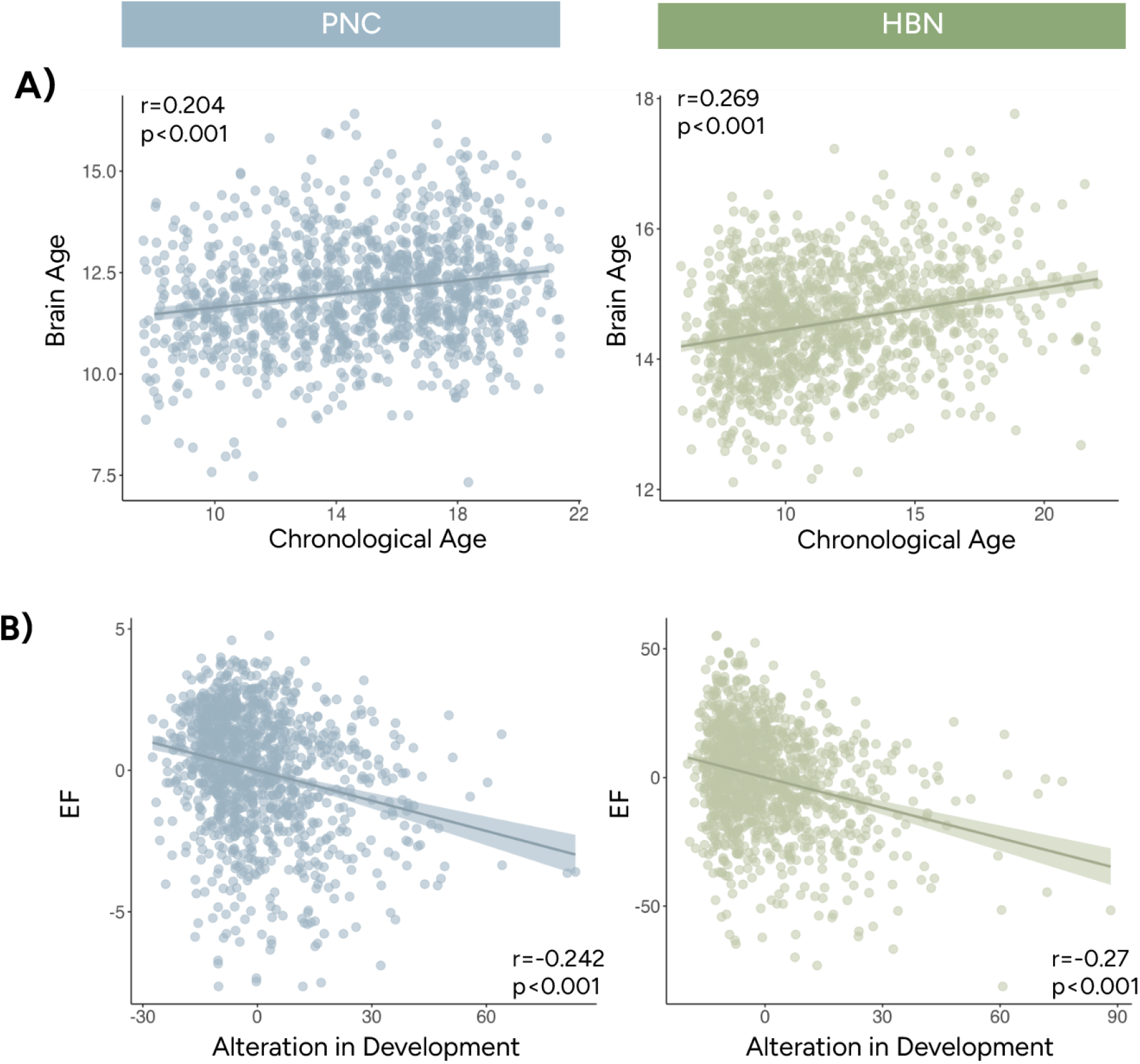
Age predictions using state engagement variability measures. **A)** Brain age predicted by state engagement variabilities was significantly associated with chronological age. **B)** Individuals with state engagement variabilities resembling someone older or younger showed worse EF performance. Age was regressed from both EF and age alteration before plotting. Instead of the actual values, residuals from the regressions were shown here to match our partial correlation analysis. Results from PNC and HBN were shown in blue and green, respectively.

### Increased brain age alteration was linked to lower EF performance

The previous analyses presented reliable evidence demonstrating that SEV changed with age and supported EF development. SEV’s robust developmental trajectory allowed us to investigate whether alterations from typical SEV development related to individual differences in EF. We measured alteration from typical development by the squared difference between chronological age and brain age estimated from SEV. Larger alterations were interpreted as appearing younger (i.e., delayed development) or older (i.e., accelerated development) compared to one’s peers regarding each individual’s SEV profile. Like the previous analysis, linear models were trained to predict age with SEV measures in the PNC and HBN rest cohorts (**Supplementary Table 9**). Brain age correlated significantly with actual age (tested in PNC: r=0.204, p<0.001; tested in HBN: r=0.269, p<0.001; Figure **6A**). Both datasets exhibited a significant negative correlation between brain age alteration and EF (tested in PNC: r=-0.242, p<0.001; tested in HBN: r=-0.27, p<0.001; Figure **6B**) when controlling for chronological age. These results indicate that both accelerated and delayed SEV development are associated with lower EF performance.

## Discussion

In over 3000 participants from four independent datasets, variability in the engagement of recurring brain states followed a nonlinear trajectory characterized by an initial stage of age-related increases before peaking in mid to late adolescence. Consistent findings generalized across resting-state, naturalistic, and task-based fMRI paradigms, suggesting that changes in SEV characterize the developing brain. Furthermore, SEV was closely associated with individual differences in EF, predicting EF performance in previously unseen individuals and partially mediating the relationship between age and EF. Furthermore, alterations from typical development of SEV were related to lower EF performance. Our results extend previous literature by supporting how neural variability allows the brain to respond dynamically to increasingly complex demands during development.

### A ubiquitous developmental trajectory observed in SEV

Our findings echo previous work reporting increased neural variability in older adolescents and young adults (4,11,14). While SEV increased with age in childhood and early adolescence, it stabilized around mid-adolescence. The initial growth of SEV can provide individuals with greater cognitive flexibility to support information processing, which is vital for increased exploration and enhanced learning about the changing world. However, future work is needed to investigate whether elevated SEV emerges naturally or serves as a response to the heightened variability in experiences during development.

The developmental trajectory of SEV demonstrated characteristics consistent with known trajectories of other brain measures. Being able to track each individual brain state continuously revealed that SEV developmental trajectory may vary by brain states. Specifically, low-cognition and cue/transition brain states reached the stabilization point before the fixation and high-cognition brain states. Low-cognition and cue/transition brain states mainly engaged lower-level or unimodal networks such as the motor and visual systems (**Supplementary Table 2**). The fixation and high-cognition brain states recruited higher-level networks in the association cortex, including the default mode and the frontoparietal networks (**Supplementary Table 2**). Brain development follows a hierarchical axis from the sensorimotor systems to the association cortex (30–33). The earlier maturation in the low-cognition and cue/transition brain states coupled with relatively later development in the fixation and high-cognition brain states dovetails with those observations.

We observed sex differences such that female participants appeared to reach the stabilization point before male participants for the fixation brain state. This finding aligns with prior evidence of sex-stratified developmental trajectories of both whole-brain and subcortical gray matter volume (19,33). The later peaks in SEV in male participants also aligned with studies demonstrating similar lags in resting-state functional connectivity stability (34), puberty emergence as well as some elements of EF performance (e.g., impulse control; 35) in male adolescents. One interesting future research direction is to probe whether sex differences in peak timing affect the emergence of disorders with significant sex differences in prevalence (36).

While the current study focused on development, future work could extend these trajectories into adulthood. Since the samples here mainly included adolescents, it is not immediately clear whether the trajectory follows an adolescent emergent (i.e., trajectory peaks at adolescence and stabilizes in adulthood; 37) or adolescent-specific trajectory (i.e., trajectory peaks in adolescence and falls in adulthood; 37). Our previous work found that SEV decreased with age in adult participants with and without psychiatric disorders (18). These preliminary results suggest that SEV might follow an adolescent-specific or inverted-U shape trajectory. However, additional research is needed to study SEV across the lifespan and its implications for EF changes in aging populations.

### SEV supports EF: a trait-like association

Here, we provided three lines of evidence demonstrating a close association between SEV and EF. SEV predicted EF and partially mediated the association between EF and age. These findings are consistent with previous studies showing a positive relationship between neural variability and faster, more accurate, and more stable performance in a range of tasks (1,4–6,38). As these associations were observed between behaviors collected outside the scanner and brain dynamic measures extracted from resting-state, naturalistic and task-based fMRI, SEV likely reflects a trait characteristic with important behavioral relevance.

A natural conclusion from these results is that increased SEV is associated with better cognition. However, the on-time development of SEV is also important, with accelerated and delayed SEV development linked to lower EF performance scores. This Goldilocks effect was observed in two independent datasets with external validation. One interesting consideration here is what contributes to alterations from typical development. Accelerated and delayed development in brain structure and functional connectivity have been associated with various risk factors, including parental deprivation, low socioeconomic status, preterm birth, childhood abuse, and stress (39–43). Factors such as social structure, community violence, and marginalization, that can be harder to measure, can also play a role in brain development (44). Delineating how SEV relates to environmental factors and how these risk factors influence neural variability remains an important future direction.

It is also important to acknowledge that alterations from typical development may be adaptive in ways not captured by the EF tasks. Such alterations may be a manifestation of an adaptive response developed in the context of adversity exposure (45,46). For instance, youth may establish accelerated SEV to cope with the demands of stress or trauma exposure. Future research will be important to better understand the function of alterations in the pace of developmental timing of SEV. In the case of accelerated development, while higher variability may increase noise and hinder the distinction between meaningful and irrelevant stimuli during an EF task, it may also facilitate more rapid responding in unpredictable contexts. Notably, recovery from stress-related functional network alterations has been observed after stress removal (47). As increased plasticity during development confers substantial potential for change (44), alterations in SEV development and their behavioral implications will likely vary as adolescents continue to be exposed to new experiences.

### Limitations and future directions

The current study provides new insight into how neural variability changes with age and supports EF, but several limitations should be considered. What causal factors underlie the change and peak in SEV is unclear. Stabilization in SEV may accompany other important biological events like puberty. Further work using longitudinal datasets with Tanner staging information is needed to better understand the observed sex differences.

Furthermore, we identified recurring brain states in the HCP dataset. It was selected for its potential to uncover brain states underpinning a range of processes, as it included paradigms spanning low-level to high-level functioning. However, as the HCP dataset mainly recruited adolescents and adult participants, it did not share much age overlap, especially with participants in the PNC and HBN datasets. Age-related changes in SEV could simply reflect adolescents’ brain states becoming more adult-like (48). If true, our results suggest that the developmental timing of when this occurs has behavioral relevance and may vary across networks. As densely sampled neuroimaging data become available, identifying brain states within each individual and extracting a personalized measure of variability may become a possibility for future research. Such studies will also allow the investigations of whether the neural variability characteristics reported here may be observed in other brain states.

In the current study, we did not explicitly exclude individuals who were at high risk or reported a clinical diagnosis. We aimed to understand SEV better by including a more diverse set of participants. Our sensitivity analysis additionally provided preliminary evidence that our result remained consistent even when clinical diagnosis was included in the model (**Supplementary Materials**). However, since many developmental and psychiatric disorders emerge during adolescence (49), it remains an interesting research direction to explore whether alterations from typical developmental trajectories of SEV might co-occur with symptom onset.

We leveraged a new computational framework to investigate variability in moment-to-moment state engagement during development. Across four independent datasets, we obtained reliable findings demonstrating that SEV changed with age. Its developmental trajectory followed the sensorimotor-to-association-cortex developmental axis and demonstrated sex differences. SEV additionally has important behavioral relevance. Notably, alterations from a typical developmental trajectory of SEV were linked to lower EF scores. These findings offer a new perspective on how variations in neural variability during development can contribute to behavioral changes.

## Methods

### Datasets and participants

This study analyzed two cross-sectional (PNC and HBN) datasets and two longitudinal (MLS and IMAGEN) neurodevelopmental datasets. PNC releases 1–2 and HBN releases 1–10 were included. Since two resting-state runs were available from HBN, we selected the one with the lower mean framewise displacement (MFD). Multiple runs of Go/No-Go were collected for MLS. However, only the first run was analyzed here to retain as many visits as possible while keeping the run number consistent across participants. For the IMAGEN cohort, we only included participants with all three visits and used all of their available Stop Signal data. No additional selection was performed for PNC rest and HBN *Despicable Me* movie data (only one run of data was collected).

### FMRI data preprocessing

The acquisition parameters for PNC, HBN, MLS, and IMAGEN have been detailed elsewhere (20,22–25,33). We applied the same standard preprocessing procedures described in previous work to the structural and functional data from all datasets (50). Structural data was nonlinearly registered to the standard MNI-152 space following brain extraction with OptiBet. We then performed slice time (in PNC and MLS only) and motion correction on the fMRI data using SPM8 before linearly aligning it to the structural data. Additional data cleaning was carried out in BioImage Suite. This included the regression of covariates of no interest, including linear and quadratic drift, a 24-parameter model of motion, and mean white matter, cerebrospinal fluid, and gray matter signals. We additionally temporally smoothed (cutoff frequency approximately at 0.12Hz) and extracted time series data using the Shen-268 atlas.

Additional quality control criteria are summarized in the Supplementary Material. Different motion thresholds were adapted to mitigate artifacts across datasets. For resting-state analysis, time points with over 0.45 framewise displacement were censored. Participants with over 20% of their time points scrubbed due to motion were excluded from further analysis. For the HBN movie, MLS Go/No-Go, and IMAGEN Stop Signal fMRI data, we avoided scrubbing to preserve the continuous nature of the paradigms. Individuals with MFD over 0.25 were excluded from the HBN movie and IMAGEN Stop Signal cohorts. Participants with MFD over 0.2 were removed from the MLS Go/NoGo cohort. Children and adolescents often showed higher levels of motion. Obtaining consistent results with different motion exclusion criteria suggests that our findings are likely to be robust to motion-induced artifacts.

After these exclusion criteria, 1208 PNC participants (female: 658; age: 14.688±3.321; age range: 8-21), 1275 HBN resting-state participants (female: 491; age: 11.693±3.39; age range: 5.96-22.07), 1312 HBN movie participants (female: 505, age: 11.361±3.606; age range: 5.13-22.07), 103 MLS cohort 1 participants (female: 40; ages 7.6-21.7 years), 150 MLS cohort 2 participants (female: 56; 16.1-28.5 years), and 530 IMAGEN participants (female: 319; data collected at 14, 19 and 22 years old) were included in our analyses.

### Variability in state engagement

A previously established approach identified a set of canonical brain states (27). Nonlinear manifold learning was used to reduce and project task-based fMRI data from the HCP into a lower-dimensional space. Time points showing similar activation patterns were located closer to each other. We then identified four brain states with distinct activation patterns using K-means clustering. These brain states reappeared across various task conditions and time points. Based on the dominant task conditions associated with time points from each brain state, we characterized them as fixation, high-cognition, low-cognition, and cue/transition. Overall, the brain networks linked to each brain state aligned with its associated cognitive processes (**Supplementary Material**).

State engagement variability was then evaluated using a recently introduced framework (18; **Figure 1**). This approach allows brain states to overlap spatially and temporally to provide a continuous, moment-to-moment measure of brain state engagement. Using this approach, the recurring brain states identified earlier were extended to PNC, HBN, MLS, and IMAGEN with non-negative least squares regression. Specifically, each brain state’s representative time point was regressed from each time point of interest from the three datasets. This step returned a beta coefficient for each state at each time point, indicating its engagement at that moment. Continuous engagement assessment allows us to examine variability in engagement for each brain state. State engagement variability was operationalized as the standard deviation of each state’s beta coefficients over time.

We used this approach to extract state engagement variability measures from all datasets analyzed here. MANOVA was used to investigate main age, sex, and age-by-sex interaction in PNC and HBN, whereas LME was used for the longitudinal MLS and IMAGEN cohorts. We included age and sex as fixed effects and participants as random effects.

### Mediation analysis with EF

To understand the behavioral implications of state engagement variability changing across development, we studied its association with EF. CNB tasks focusing on the complex cognition and executive control domains were analyzed here (see **Supplementary Material** for the specific tasks included). An efficiency score was computed for each task based on previous literature (51). Response and accuracy scores were first standardized across participants. We then multiplied the response score by −1 before summing it with the accuracy score. For HBN, the standardized scores from the Card Sort, Flanker, List Sorting Working Memory, and Pattern Comparison Process Speed paradigms were used. After removing individuals with missing behavioral data, 1161 PNC, 1189 HBN rest, and 1229 HBN movie participants were included in the EF and state engagement variability analysis.

We performed PCA on the EF scores within each cohort to extract a summary measure. An additional PCA was run on the four state engagement variability measures to estimate overall state engagement variability. We then utilized partial correlation analysis to investigate whether EF was associated with overall state engagement variability, controlling for age. The mediation model (implemented with the Mediation Toolbox; 28) explored whether overall state engagement variability mediated the changes in EF with age.

## Acknowledgement

This work received support from the following sources: Gruber Science Fellowship; the European Union-funded FP6 Integrated Project IMAGEN (Reinforcement-related behaviour in normal brain function and psychopathology) (LSHM-CT-2007-037286), the Horizon 2020 funded ERC Advanced Grant ‘STRATIFY’ (Brain network based stratification of reinforcement-related disorders) (695313), Horizon Europe ‘environMENTAL’, grant no: 101057429, UK Research and Innovation (UKRI) Horizon Europe funding guarantee (10041392 and 10038599), Human Brain Project (HBP SGA 2, 785907, and HBP SGA 3, 945539), the Chinese government via the Ministry of Science and Technology (MOST). The German Center for Mental Health (DZPG), the Bundesministerium für Bildung und Forschung (BMBF grants 01GS08152; 01EV0711; Forschungsnetz AERIAL 01EE1406A, 01EE1406B; Forschungsnetz IMAC-Mind 01GL1745B), the Deutsche Forschungsgemeinschaft (DFG project numbers 186318919 [FOR 1617], 178833530 [SFB 940], 386691645 [NE 1383/14-1], 402170461 [TRR 265], 454245598 [IRTG 2773]), the Medical Research Foundation and Medical Research Council (grants MR/R00465X/1 and MR/S020306/1), the National Institutes of Health (NIH) funded ENIGMA-grants 5U54EB020403-05, 1R56AG058854-01 and U54 EB020403 as well as NIH R01DA049238, the National Institutes of Health, Science Foundation Ireland (16/ERCD/3797). NSFC grant 82150710554. Further support was provided by grants from: – the ANR (ANR-12-SAMA-0004, AAPG2019 – GeBra), the Eranet Neuron (AF12-NEUR0008-01 – WM2NA; and ANR-18-NEUR00002-01 – ADORe), the Fondation de France (00081242), the Fondation pour la Recherche Médicale (DPA20140629802), the Mission Interministérielle de Lutte-contre-les-Drogues-et-les-Conduites-Addictives (MILDECA), the Assistance-Publique-Hôpitaux-de-Paris and INSERM (interface grant), Paris Sud University IDEX 2012, the Fondation de l’Avenir (grant AP-RM-17-013), the Fédération pour la Recherche sur le Cerveau. Data were provided in part by the WU-Minn Human Connectome Project Consortium (principal investigators, David Van Essen and Kamil Ugurbil; Grant No. 1U54MH091657 funded by the 16 NIH institutes and centers that support the NIH Blueprint for Neuroscience Research; also funded by the McDonnell Center for Systems Neuroscience at Washington University) and CNP (NIH Roadmap for Medical Research Grant Nos. UL1-DE019580, RL1MH083268, RL1MH083269, RL1DA024853, RL1MH083270, RL1LM009833, PL1MH083271, and PL1NS062410).

