## Supplementary Materials for "Variation in moment-to-moment brain state engagement changes across development and contributes to individual differences in executive function"

**Brain states information:**

While there are several ways to identify recurring brain states, we replicated our prior work here (1,2). In brief, we applied nonlinear manifold learning and 2-step Diffusion Mapping to project task-based fMRI data from the Human Connectome Project S500 release (3) into a lower-dimensional space. Data from the minimal HCP preprocessing pipeline was used here. We included six tasks (motor, working memory, social, emotional, relational, and gambling) from 390 participants for state identification after quality control (1). As this dataset covers a range of paradigms, it has the potential to uncover brain states underlying different cognitive processes. Identifying brain states in an external dataset also helps mitigate potential overfitting concerns.

After fMRI data was reduced and projected into a lower-dimensional space, time points showing similar activation patterns were closer. K-means clustering was then used to identify 4 recurring brain states with distinct activation patterns. The number of brain states was determined using the Calinski-Harabasz criterion (4). Based on the dominant task conditions in each brain state, we characterized them as fixation, high-cognition, low-cognition, and cue/transition (**Supplementary Table 1**). We identified the centroid of each state cluster to serve as its representative time point for later analyses.

Our previous work also investigated the brain networks involved in these brain states. For each representative time point, we first identified the activated and deactivated brain regions in a set of canonical brain networks (defined as having activation above or below 0, respectively). The activation or deactivation percentage was computed by dividing the number of activated or deactivated brain regions by the total number of brain regions in a network.

We found that different brain networks were activated differently in each brain state (**Supplementary Table 2**). For instance, the visual networks showed high activation percentages in the high-cognition brain state but low activation during fixation. This may be partly due to participants being shown more complex visual stimuli during tasks than fixation. The activation of different brain networks also followed what cognitive processes were associated with each state. For example, the entire motor network was activated for the low-cognition state, which was strongly associated with the motor task.

**Supplementary Table 1.** Number of volumes associated with each task condition for the four recurring brain states

| | Fixation | High-cognition | Low-cognition | Cue/Transition |
|---|---|---|---|---|
| Fixation | 635 | 0 | 20 | 65 |
| Cue | 41 | 3 | 6 | 158 |
| Working memory (0 back) | 10 | 56 | 99 | 123 |
| Working memory (2 back) | 1 | 201 | 10 | 76 |
| Emotion (Fear) | 10 | 42 | 0 | 48 |
| Emotion (Neutral) | 23 | 12 | 99 | 16 |
| Gambling (Win) | 0 | 100 | 25 | 35 |
| Gambling (Loss) | 0 | 101 | 10 | 49 |
| Motor (Tongue) | 0 | 1 | 52 | 12 |
| Motor (Left foot) | 8 | 5 | 41 | 13 |
| Motor (Left hand) | 7 | 0 | 45 | 15 |
| Motor (Right foot) | 0 | 0 | 55 | 12 |
| Motor (Right hand) | 0 | 1 | 50 | 15 |
| Social (Mental) | 0 | 113 | 0 | 47 |
| Social (Random) | 0 | 111 | 0 | 109 |
| Relational (Match) | 3 | 9 | 17 | 40 |
| Relational (Relation) | 3 | 90 | 5 | 9 |

**Supplementary Table 2.** Networks showing the highest activation and deactivation percentages for each state

| | Fixation | High-cognition | Low-cognition | Cue/transition |
|---|---|---|---|---|
| Activation percentages | DMN (88.89%) | VAs (100%) | Motor network (100%) | Visual I (100%) |
| | Motor network (85.71%) | Visual II (88.87%) | MF (86.21%) | VAs (72.22%) |
| | MF (82.76%) | FP (82.35%) | Cerebellum (84%) | Visual II (66.67%) |
| Deactivation percentages | VAs (94.44%) | Motor network (87.76%) | Visual I (100%) | MF (89.66%) |
| | Visual I (66.67%) | DMN (83.33%) | Visual II, VAs, and DMN (66.67%) | DMN (88.87%) |
| | Visual II (66.67%) | Subcortical (79.31%) | | Motor (83.67%) |

This table shows the canonical functional networks with the three highest activation and deactivation percentages for each brain state. The actual activation and deactivation percentage values were included in parentheses. DMN, default mode network; MF, medial frontal network; VAs, visual association network; FP, frontoparietal network.

**Functional magnetic resonance imaging preprocessing and quality control:**

For the PNC rest cohort, we retained 1441 participants after excluding individuals failing preprocessing benchmarks (i.e., missing resting-state fMRI data, issues with skull stripping, nonlinear or linear registration). All participants had the same number of volumes (N=124). We removed individuals if they were missing brain coverage (N=27), had over 20% of their volumes scrubbed due to motion (N=72), or were missing age information (N=17). To ensure that no one time point showed a significantly worse fit than the others during non-negative least squares regression, we removed participants if any of their time points showed a significant outlier residual (N=1).

For the HBN rest cohort, 1588 participants remained after excluding individuals who did not pass the preprocessing benchmarks or had missing resting-state data. We also removed individuals with an incomplete scan (fewer than 375 volumes; N=48). Similar to the PNC rest cohort, individuals with missing brain coverage (N=148), over 20% of their volumes scrubbed due to motion (N=38), and missing age information (N=79) were excluded from further analysis. No participant showed a significant outlier residual in the model.

For the HBN movie cohort, we had 1718 participants after excluding people who did not meet preprocessing benchmarks. To preserve the continuous nature of the naturalistic paradigm, we did not censor volumes based on motion here. Instead, individuals with mean framewise displacement (MFD) over 0.25mm were excluded (N=195). We additionally removed participants with an incomplete scan (fewer than 750 volumes; N=49), missing brain nodes (N=80), or behavioral information (N=82). No participant in this cohort had an outlier residual for any of their time points.

Preprocessing for the MLS longitudinal data was performed slightly differently. Exclusion criteria were first applied to each visit. We retained 493 cohort 1 visits and 669 cohort 2 visits after selecting ones that passed all preprocessing benchmarks and had a MFD less than 0.2mm for the first Go/No-Go task run. Every task run had 93 volumes. To retain as many visits as possible for longitudinal analysis, we removed brain nodes with incomplete coverage from all participants (instead of runs with missing brain coverage, 16 and 18 brain nodes were removed from cohorts 1 and 2, respectively). Four visits from cohort 2 were excluded due to having an outlier residual during regression. The last exclusion criterion was at the participant level. We removed participants with only 1 available visit (final N: cohort 1: 479 visits, 103 unique participants; cohort 2: 639 visits, 150 unique participants; see **Supplementary Figure 1** for further visit breakdown).

To visualize the longitudinal trajectories and explore potential sex differences, we combined both MLS cohorts together. For consistency, we re-extracted state engagement variability measures after removing the same set of brain nodes with missing coverage (N=19) from individuals in both cohorts. Additionally, one cohort 1 and four cohort 2 participants were excluded due to having outlier residuals here. The final analysis included 1116 visits and 252 unique participants.

For the IMAGEN dataset, we only analyzed participants with all three available visits. After preprocessing exclusion criteria were applied, we retained 660 participants. We excluded individuals if their visits showed MFD over 0.25 (N=116). Almost all participants had the same number of volumes from visit 1 (444 volumes). Still, the volume numbers varied in visit 2 (most participants had between 300–350 volumes) and 3 (most participants had between 300–370 volumes). For this reason, we decided to exclude anyone if they had fewer than 444 volumes for visit 1 (N=2) or fewer than 300 volumes for visit 2 and 3 (N=2). We additionally excluded participants with missing brain coverage (N=4). Four additional participants were removed from all analyses due to having outlier residuals. After all exclusion criteria, we included 530 participants with both brain and age information for our analysis.

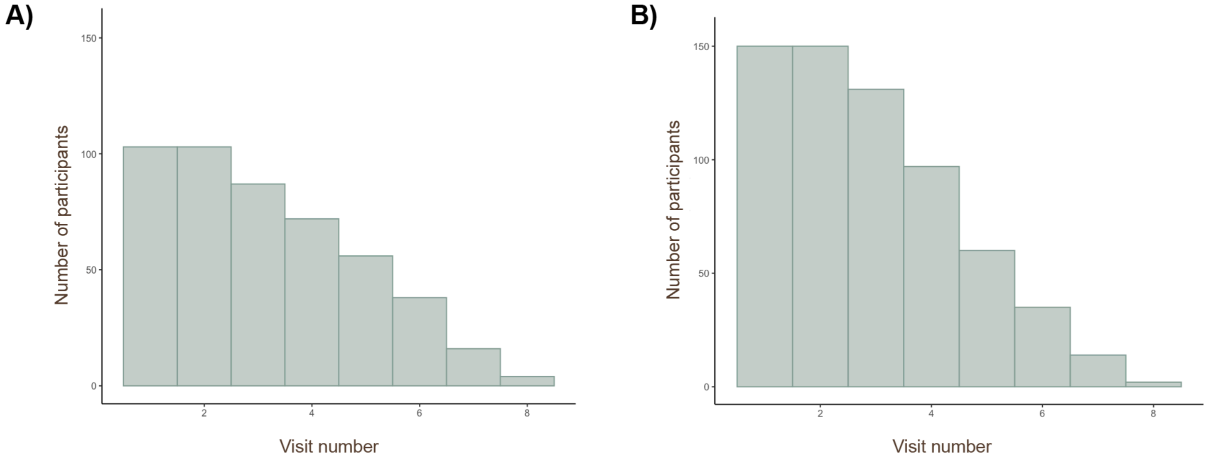


**Supplementary Figure 1.** Visit number for MLS. The two histograms showed how many participants were included for each visit number for MLS cohort 1 **A)** and cohort 2 **B)**.

**PNC EF measures**

From the PNC CNP, we selected tasks measuring the executive control and complex cognition domains based on previous work (5). The Penn Conditional Exclusion Test, Penn Continuous Performance Test, and Penn Letter N-back Test evaluated executive control, whereas the Penn Verbal Reasoning Test, Penn Matrix Reasoning Test, and Penn Line Orientation Test assessed complex cognition.

**Supplementary Table 3.** Visit effects on state engagement variability in IMAGEN.

| Fixation | | | |
|---|---|---|---|
| Predictor | Beta | T-value | P-value |
| Age 14 vs. Age 19 | 2.431e-02 | t(1056)=2.451 | 0.014 |
| Age 14 vs. Age 22 | 3.736e-02 | t(1056)=3.767 | <0.001 |
| Age 19 vs. Age 22 | 1.305e-02 | t(1056)=1.316 | 0.189 |
| High-cognition | | | |
| Predictor | Beta | T-value | P-value |
| Age 14 vs. Age 19 | 1.374e-02 | t(1056)=2.101 | 0.036 |
| Age 14 vs. Age 22 | 1.507e-02 | t(1056)=2.304 | 0.021 |
| Age 19 vs. Age 22 | 1.325e-03 | t(1056)=0.203 | 0.840 |
| Low-cognition | | | |
| Predictor | Beta | T-value | P-value |
| Age 14 vs. Age 19 | -7.699e-04 | t(1056)=-0.403 | 0.687 |
| Age 14 vs. Age 22 | -1.190e-03 | t(1056)=-0.622 | 0.534 |
| Age 19 vs. Age 22 | -4.203e-04 | t(1056)=-0.220 | 0.826 |
| Cue/transition | | | |
| Predictor | Beta | T-value | P-value |
| Age 14 vs. Age 19 | 1.092e-02 | t(1056)=1.767 | 0.078 |
| Age 14 vs. Age 22 | 7.844e-03 | t(1056)=1.269 | 0.205 |
| Age 19 vs. Age 22 | -3.077e-03 | t(1056)=-0.498 | 0.619 |

**Supplementary Table 4. LME other covariates for MLS cohort 1**

| Fixation | | | |
|---|---|---|---|
| Predictor | Beta | T-value | P-value |
| Sex | 0.036 | 0.380 | 0.704 |
| Age-by-sex | -0.004 | -0.654 | 0.513 |
| High-cognition | | | |
| Predictor | Beta | T-value | P-value |
| Sex | -1.217e-02 | -0.199 | 0.843 |
| Age-by-sex | -3.637e-04 | -0.083 | 0.934 |
| Low-cognition | | | |
| Predictor | Beta | T-value | P-value |
| Sex | -1.360e-02 | -0.811 | 0.418 |
| Age-by-sex | 4.314e-04 | 0.360 | 0.719 |
| Cue/transition | | | |
| Predictor | Beta | T-value | P-value |
| Sex | 0.029 | 0.516 | 0.606 |
| Age-by-sex | -0.003 | -0.738 | 0.461 |

**Supplementary Table 5. LME other covariates for MLS cohort 2**

| Fixation | | | |
|---|---|---|---|
| Predictor | Beta | T-value | P-value |
| Sex | 0.123 | 0.801 | 0.424 |
| Age-by-sex | -0.005 | -0.745 | 0.457 |
| High-cognition | | | |
| Predictor | Beta | T-value | P-value |
| Sex | 4.682e-02 | 0.483 | 0.629 |
| Age-by-sex | -1.925e-03 | -0.453 | 0.651 |
| Low-cognition | | | |

| Predictor | Beta | T-value | P-value |
| --- | --- | --- | --- |
| Sex | -6.277e-03 | -0.235 | 0.814 |
| Age-by-sex | 3.064e-04 | 0.261 | 0.794 |
| Cue/transition | | | |
| Predictor | Beta | T-value | P-value |
| Sex | 0.068 | 0.711 | 0.477 |
| Age-by-sex | -0.002 | -0.557 | 0.577 |

**Supplementary Table 6. MANOVA other covariates (PNC and HBN)**

| PNC Rest | | |
|---|---|---|
| Predictor | F-stat | P-value |
| Sex | F(4,1201)=1.133 | 0.340 |
| HBN Rest | | |
| Predictor | F-stat | P-value |
| Sex | F(4,1268)=1.884 | 0.111 |
| HBN Movie | | |
| Predictor | F-stat | P-value |
| Sex | F(4,1305)=4.084 | 0.003 |

**Supplementary Table 7. LME other covariate for IMAGEN**

| Fixation | | | |
|---|---|---|---|
| Predictor | Beta | T-value | P-value |
| Sex | 1.916e-02 | t(1315)=1.422 | 0.155 |
| (Difference between Age 14 and 19)-by-sex | 2.527e-02 | t(1056)=1.607 | 0.108 |
| (Difference between Age 19 and 22)-by-sex | 2.863e-02 | t(1056)=1.821 | 0.069 |
| High-cognition | | | |
| Predictor | Beta | T-value | P-value |
| Sex | 1.496e-03 | t(1269)=0.164 | 0.870 |
| (Difference between Age 14 and 19)-by-sex | 2.042e-02 | t(1056)=1.970 | 0.049 |
| (Difference between Age 19 and 22)-by-sex | 1.699e-02 | t(1056)=1.639 | 0.102 |
| Low-cognition | | | |
| Predictor | Beta | T-value | P-value |
| Sex | -2.606e-04 | t(1322)=-0.101 | 0.920 |
| (Difference between Age 14 and 19)-by-sex | 6.861e-03 | t(1056)=2.264 | 0.024 |
| (Difference between Age 19 and 22)-by-sex | 5.044e-03 | t(1056)=1.664 | 0.096 |
| Cue/transition | | | |
| Predictor | Beta | T-value | P-value |
| Sex | 2.481e-02 | t(1274)=2.890 | 0.004 |
| (Difference between Age 14 and 19)-by-sex | 8.386e-03 | t(1056)=0.856 | 0.392 |
| (Difference between Age 19 and 22)-by-sex | 2.586e-02 | t(1056)=2.640 | 0.008 |

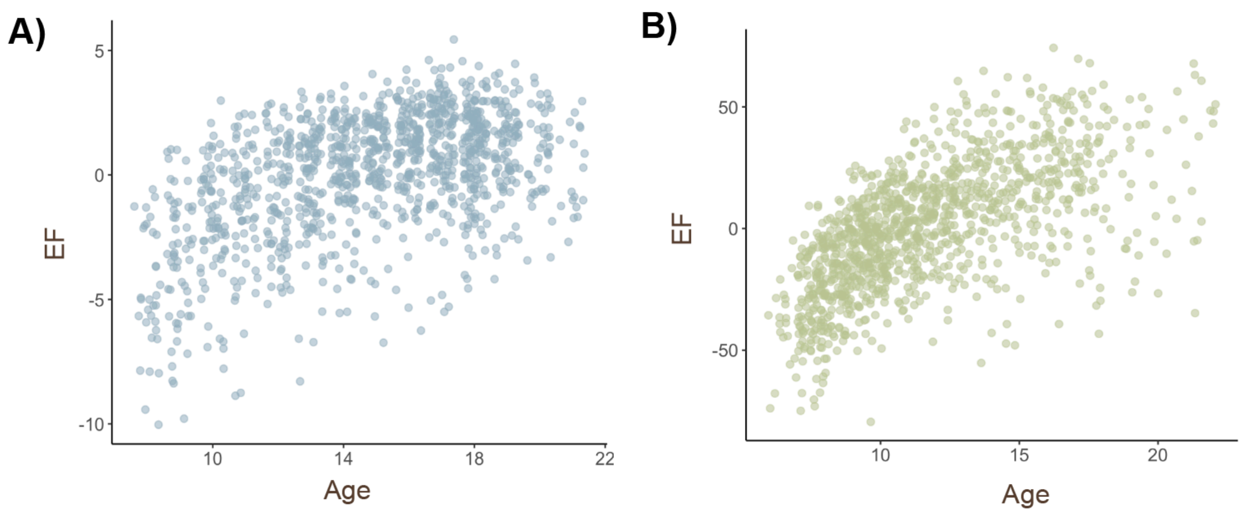


**Supplementary Figure 2**. EF by age in PNC **A)** and HBN **B)**.

**Supplementary Table 8. Model parameters from predicting EF using state engagement variabilities**

| Brain states/Model coefficients | Trained in HBN/Tested in PNC | Trained in PNC/Tested in HBN |
|---|---|---|
| Fixation | -20.203 | 1.628 |
| High-cognition | 83.631 | 2.671 |
| Low-cognition | -5.087 | 6.675 |
| Cue/transition | 6.951 | -4.231 |

**Supplementary Table 9. Model parameters from predicting age using state engagement variabilities**

| Brain states/Model coefficients | Trained in HBN/Tested in PNC | Trained in PNC/Tested in HBN |
|---|---|---|
| Fixation | 0.412 | 0.518 |
| High-cognition | 11.027 | 10.112 |
| Low-cognition | 3.679 | 8.032 |
| Cue/transition | -2.780 | -8.733 |

**Note:** Models were first trained using all participants with available age and state engagement variability information before being applied to unseen individuals with available behavior and state engagement variability data.

Sensitivity analysis revealed consistent results when controlling for clinical diagnosis

As we did not exclude participants diagnosed with psychiatric disorders, we further examined whether results would remain consistent after controlling for diagnostic status. We mainly focus on the HBN dataset here, as the majority of HBN participants were diagnosed with psychiatric disorders. Participants were separated into two groups based on whether they had received a clinical diagnosis or not. We reran our MANOVA analyses using only participants with available diagnostic information (rest: 69 participants with no diagnosis, 831 participants with at least one diagnosis; movie: 87 participants with no diagnosis, 850 with at least one diagnosis). As in the primary findings, state engagement variability increased with age when including clinical diagnosis as a covariate (rest: $F(4,892)=31.266$, $p<0.001$; movie: $F(4,926)=47.967$, $p<0.001$). Clinical diagnosis did not show a main effect on state engagement variability (rest: $F(4,892)=0.409$, $p=0.803$; movie: $F(4,926)=1.957$, $p=0.099$). Alterations from typical state engagement variability were linked to worse EF performance when controlling for age and clinical diagnosis ($r=-0.267$; $p<0.001$). These results suggest that the observed developmental patterns remain robust even after accounting for the diagnostic group.
